# Decoupling of the nuclear division cycle and cell size control in the coenocytic cycle of the ichthyosporean *Sphaeroforma arctica*

**DOI:** 10.1101/190900

**Authors:** Andrej Ondracka, Iñaki Ruiz-Trillo

**Affiliations:** Institut de Biologia Evolutiva (CSIC-Universitat Pompeu Fabra), Passeig Maritím de la Barceloneta 37-49, 08003 Barcelona, Spain; Departament de Genètica, Microbiologia i Estadística, Universitat de Barcelona, Barcelona, Catalonia, Spain.; ICREA, Pg. Lluís Companys 23, 08010 Barcelona.

## Abstract

Coenocytes (multinucleated cells formed by sequential nuclear divisions without cytokinesis) are commonly found across the eukaryotic kingdom, including in animals, plants and several lineages of unicellular eukaryotes. Despite their commonality, little is known about how cell growth, nuclear divisions and cell divisions are coordinated in coenocytes. Among the unicellular eukaryotes that form coenocytes are ichthyosporeans, a lineage of unicellular holozoans that are of significant interest due to their phylogenetic placement as one of the closest relatives to animals. Here, we characterize the coenocytic cell division cycle in the ichthyosporean *Sphaeroforma arctica*. In laboratory conditions, we observed that *S. arctica* cells undergo a highly regular periodic coenocytic cell cycle. Nuclear division cycles occur synchronously within the coenocyte and in regular time intervals (~11 hours per nuclear cycle) until reaching 64-128 nuclei and releasing daughter cells. The duration of the nuclear division cycles is constant across a wide range of nutrient concentration. In contrast, the volume of the coenocytes increase more slowly in lower nutrient concentration, which also results in smaller newborn daughter cells. This suggests that *S. arctica* cells are capable to adapt the cell growth rate to nutrient concentration while maintaining the timing of nuclear division cycles, suggesting that in ichthyosporeans the mechanisms regulating highly periodic nuclear division cycles operate independently from mechanisms sensing the cell size.

## Introduction

To adapt to environmental conditions, cells are capable of adapting their growth rates. Thus, the growth rate of a cell is dependent on a variety of environmental factors, including temperature and the type and concentration of the carbon or nitrogen source (Monod, 1949; Schmidt-Glenewinkel & Barkai, 2014; Ziv, Siegal, & Gresham, 2013). Moreover, cell growth can be summarized by both the rate of increase of cell size (cell volume), and the frequency of cell division. Only when these two parameters are coupled, the cell size is maintained constant over generations. Thus, to achieve balanced growth and maintain constant cell size, cells must be able to sense their size and coordinate cell growth and cell division (Amodeo & Skotheim, 2015). Such mechanisms of size sensing and the coupling of cell size to cell division have been observed across diverse eukaryotic taxa (Turner, Ewald, & Skotheim, 2012; Tzur, Kafri, LeBleu, Lahav, & Kirschner, 2009; Umen, 2005).

The prevailing view is that in unicellular eukaryotes that proliferate through mitotic cell cycle the cell size control is executed by cell size checkpoints. Those checkpoints prevent the cells to continue the cell division cycle before reaching certain size threshold. Cell size checkpoints were initially identified in budding (Johnston, Pringle, & Hartwell, 1977) and fission yeasts (Fantes & Nurse, 1977). Cell size control ensures that in poor nutrient conditions, when cell division cycles become longer, cells divide at approximately the same size regardless of the duration of the cell cycle (Brauer et al., 2008; Di Talia, Skotheim, Bean, Siggia, & Cross, 2007; Jorgensen, Nishikawa, Breitkreutz, & Tyers, 2002). A cell size checkpoint has also been described in the unicellular alga *Chlamydomonas reinhardtii* (Fang, De Los Reyes, & Umen, 2006), which divides by a multiple fission mechanism and can produce a variable number of daughter cells in each cycle. Cell size control in *Chlamydomonas* ensures that the daughter cells are all born at similar size, regardless of the size of the mother cell before divisions (Craigie & Cavalier-Smith, 1982; Donnan & John, 1983).

In contrast to unicellular organisms, cell cycle in early animal embryos is driven by a timer that ensures rapid, precisely timed and synchronous divisions (O’Farrell, Stumpff, & Su, 2004). Cell divisions in animal embryos occur in constant volume and therefore do not need to be coupled to cell growth as in unicellular organisms. A particularly well-studied system for early embryonic cell cycles is the Drosophila syncytial pre-blastoderm and blastoderm. The first rounds of nuclear divisions in the syncytium upon fertilization proceed synchronously and in regular time intervals (Farrell & O’Farrell, 2014).

The relationship between cell division cycle and cell growth can become more complicated in unicellular organisms that exhibit more complex and elaborate life cycles than, for example, the budding or fission yeast. A particular example of such complex cycle is the coenocytic cell cycle, in which cells undergo multiple nuclear divisions without accompanying cytokinesis, resulting in a single cell (coenocyte) containing multiple nuclei. Coenocytes are commonly found in a variety of unicellular eukaryotic taxa, such as filamentous fungi and several groups of protists (Adl et al., 2005), as well as in animal tissues and plants. However, with the exception of the filamentous fungus *Ashbya gossypii*, where multiple nuclei within the shared cytoplasm behave autonomously and proceed through the nuclear cycles independently of each other (Anderson et al., 2013; Dundon et al., 2016), little is known about how cell growth, nuclear divisions and cell divisions are coordinated in unicellular organisms that proliferate through a coenocytic cycle.

Here, we investigate the coupling of cell size and cell cycle progression in the coenocytic development of ichthyosporeans, a lineage of unicellular holozoans that is of interest due to its phylogenetic placement as one of the closest unicellular relatives to animals (figure 1A). Among ichthyosporeans, several species have been described to form multinucleated coenocytes (A. de Mendoza, Suga, Permanyer, Irimia, & Ruiz-Trillo, 2015; Glockling, Marshall, & Gleason, 2013; Marshall, Celio, McLaughlin, & Berbee, 2008; L. Mendoza, Taylor, & Ajello, 2002; Suga & Ruiz-Trillo, 2013), and it has even been suggested that there might be a direct evolutionary relationship between ichthyosporean coenocytes and animal multicellularity (Suga & Ruiz-Trillo, 2013). In this work, we focus on *Sphaeroforma arctica*, an ichthyosporean first isolated from an arctic marine amphipod (Jøstensen, Sperstad, Johansen, & Landfald, 2002) and whose genome has been sequenced (Grau-Bové et al., 2017). Due to its rather simple, linear life cycle (see below; (Jøstensen et al., 2002), *S. arctica* is an attractive model to study the coenocytic cell cycle of unicellular eukaryotes.

**Figure 1:**
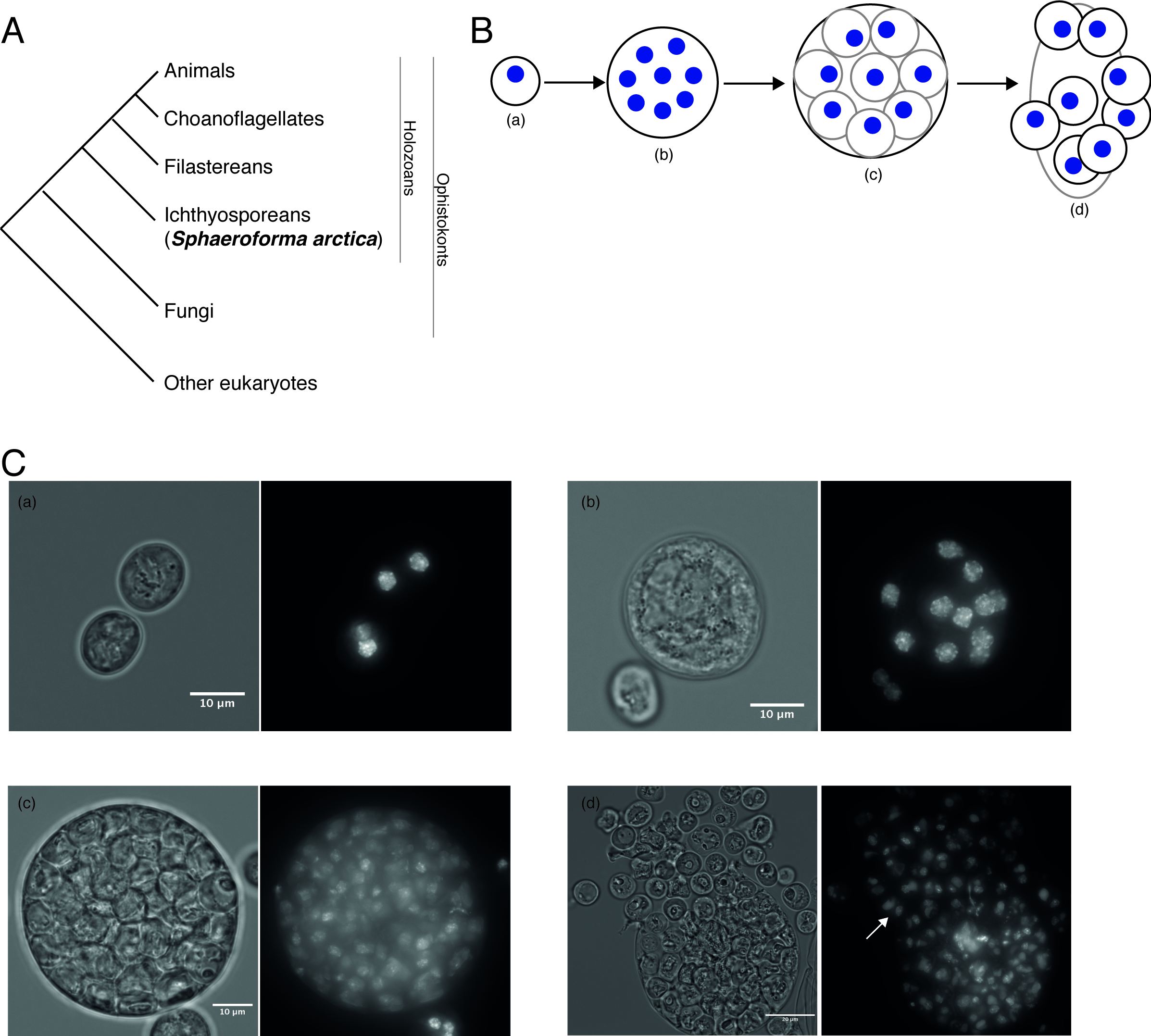
A) A cladogram representing the position of *S. arctica* within eukaryotes as in (Torruella et al., 2015) B) A schematic illustration of the *S. arctica* cell cycle comprising mono‐ or di-nucleated newborn cells (a), multinuclear coenocyte (b), cellularized coenocyte (c), and burst (d). Blue spots represent nuclei. C) Representative DIC (left) and DAPI (right) images of cells from the corresponding cell cycle stage (newborn (a), multinuclear coenocyte (b), cellularized coenocyte (c), and burst (d)). White arrow represents a newborn cell with two nuclei. Scale bar in (a-c) 10 um, in (d) 20 um.

## Results and discussion

### Cell cycle progression

We first characterized the life cycle of *S. arctica* in laboratory conditions by microscopy. *S. arctica* cultures grow at 12°C in Difco marine broth (MB) medium. The cells in the culture vary in size and number of nuclei, but exhibit uniformly round morphology (fig 1C), which could suggest a simple, linear coenocytic life cycle (fig 1B). Small, newborn cells grow into a multinucleate coenocyte by rounds of synchronous nuclear divisions (Suga & Ruiz-Trillo, 2013). This is followed by cellularization and release (burst) of the daughter cells. We observed that newborn cells frequently contain 2 or even 4 nuclei (figure 1C, panel d), white arrow). This suggests that nuclear divisions already occur inside the cellularized coenocyte before the burst, or that cellularization can occur around multiple nuclei.

We also observed that saturated cultures (grown for >7 days after inoculation into fresh media) contain almost exclusively newborn cells with low DNA content (corresponding to 1, 2 or 4C DNA content, figure 2A, time 0). This enabled us to easily synchronize the cells by starvation and examine the progression throughout the coenocytic cycle by DAPI staining for DNA content using flow cytometry. The observed peaks in DNA content profiles corresponded to two-fold increases in fluorescence intensities (figure 2A), consistent with previous findings that nuclear divisions within the coenocyte are synchronized (Suga & Ruiz-Trillo, 2013), and suggesting that DNA replication also occurs synchronously within nuclei of the coenocyte.

**Figure 2:**
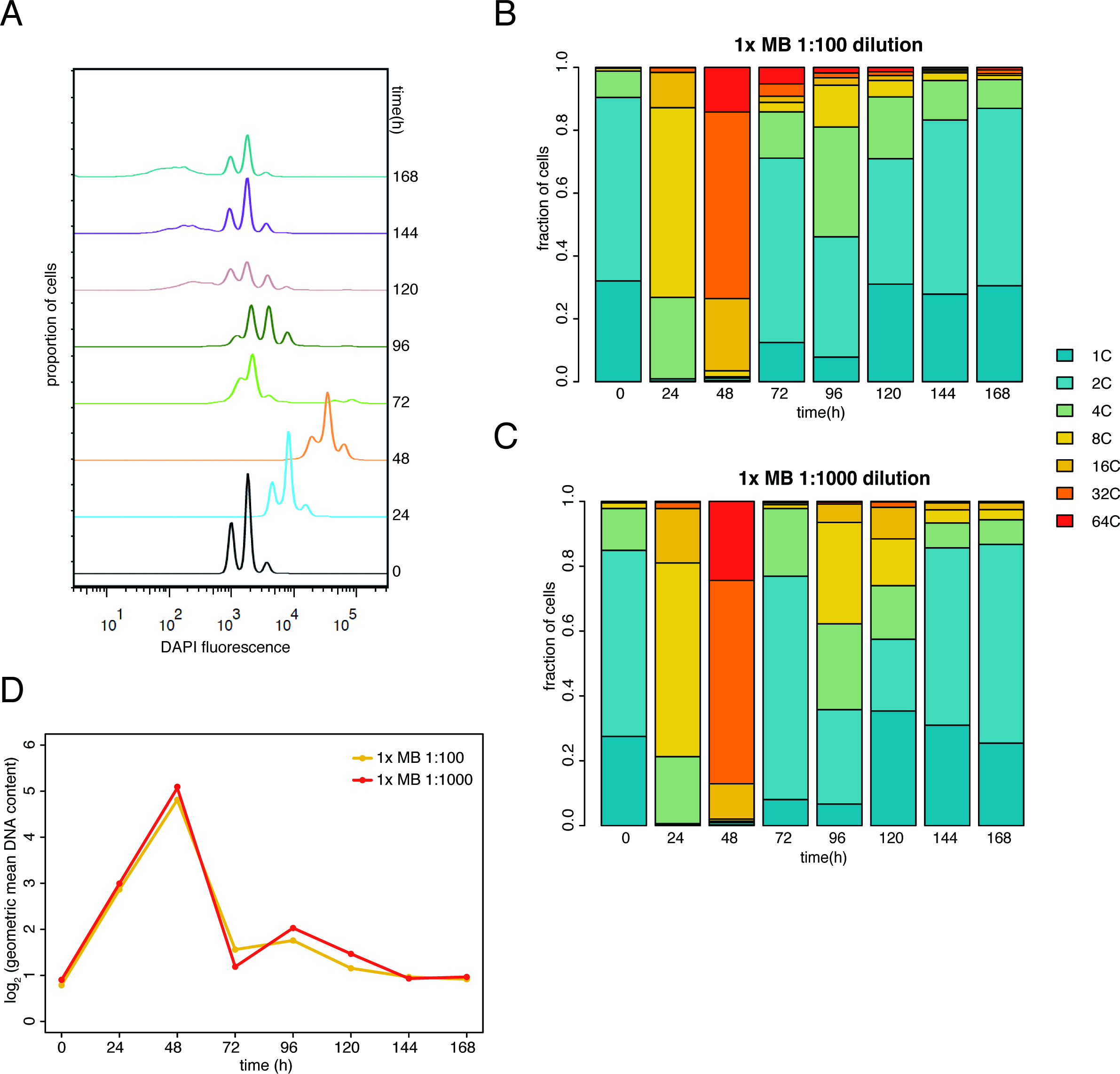
A) DNA content profile assessed by flow cytometry across the time course of cell populations grown in standard conditions (1x Marine Broth, 12°C, 1:100 initial dilution of a saturated culture. B) Quantification of fractions of population per DNA content profiles bin. C) Same as B for 1:1000 initial dilution of a saturated culture. D) Quantification of mean nuclear content at each time point (expressed as log_2_ of geometric mean).

To quantify the fraction of populations of each nuclear content, we co-stained multiple samples containing cells of different stages of the coenocytic cycle, use these bins to calibrate the DNA content based on the lowest intensity peak observed (supplementary figure 1B) and quantified the populations into bins with discrete nuclear content values (figure 2B). The results show that cells progressed through nuclear division cycles with high synchrony (all cells in the population increased DNA content at a similar rate). Cells underwent approximately 4-fold increase in DNA content per 24 hours (figure 2B), corresponding to two synchronous nuclear division cycles (rounds of DNA replication and mitosis). Between 48 and 72 hours, the majority of coenocytes burst and gave rise to newborn daughter cells. We observed that the timing of nuclear divisions, timing of burst and nuclear content of coenocytes at burst were independent of the initial dilution of the culture (figures 2B,C), indicating that cell density and remaining concentration of nutrients in the media has no effect on the progression through the coenocytic cycle.

Next, we further characterized the syncytial cell cycles under different experimental conditions. First, we tested the effect of temperature on the timing of nuclear divisions. We observed that higher temperatures speed up the rate of nuclear divisions (supplementary figure 2A-C), although the nuclear content of the coenocyte at burst and final density of cells at saturation remained the same regardless of temperature (supplementary figure 2D). Thus, temperature has an effect on the rate of nuclear division.

We then inquired whether nutrient concentration has any effect on nuclear division cycles. To do that, we carried out growth experiments with a series of media prepared by dilution of the MB medium with artificial seawater. In order to maintain approximately constant nutrient concentration during the first coenocytic cycle, the saturated cultures were heavily diluted (1:1000 initial culture dilution). We observed that the cells progressed through the cycle synchronously under all nutrient concentrations (figure 3A-D). The rate of nuclear content increase was constant throughout the coenocytic growth for all conditions (figure 3E). Although cells in lower media concentrations bursted earlier and at slightly lower nuclear content (presumably due to consumption of nutrients; figure 3A-E), the doubling time of nuclear content (approximately 11 hours) was constant and independent of nutrient concentration across the wide range (1x to 1/16x) of media concentrations (figure 3E,F).

**Figure 3:**
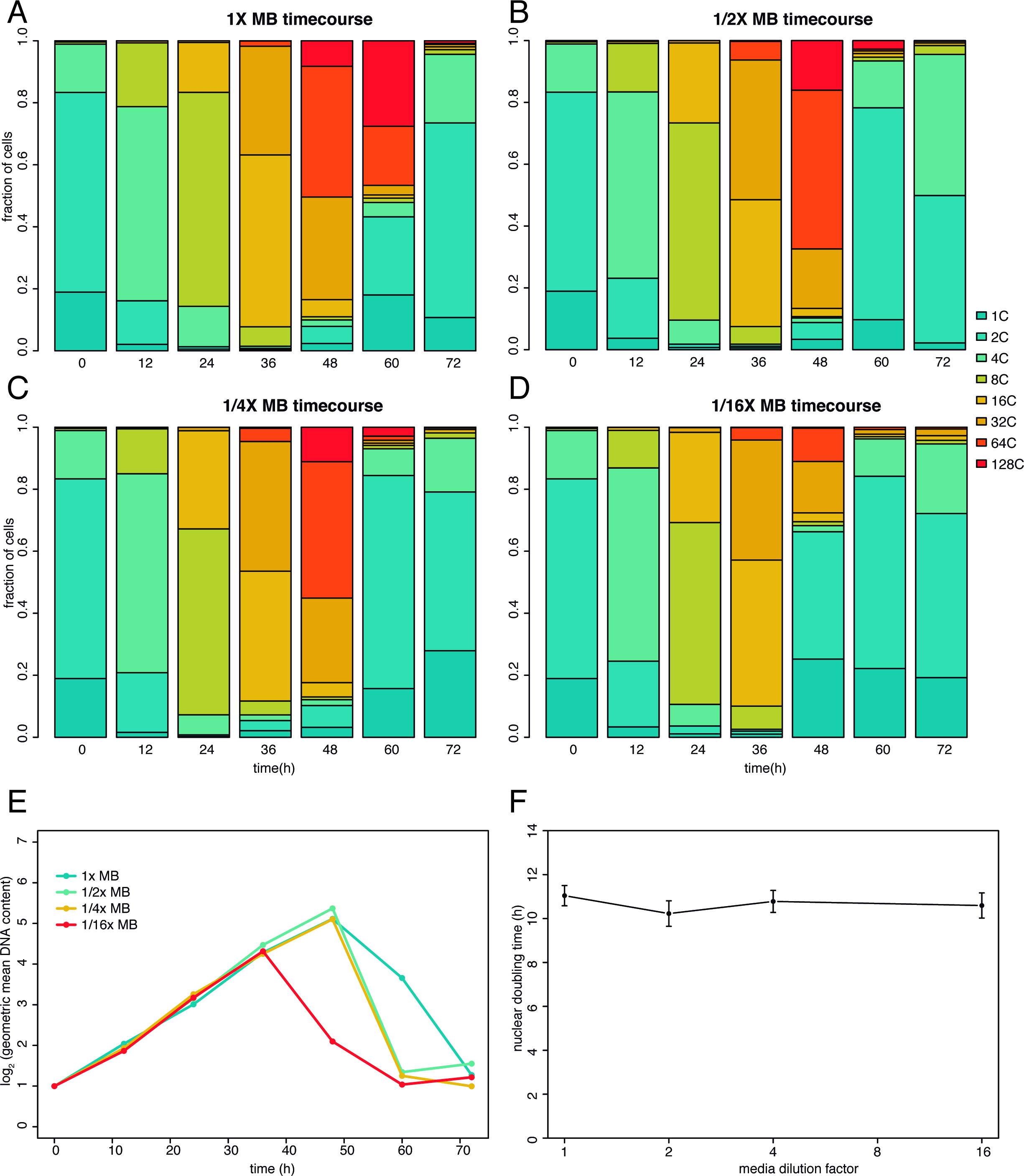
A-D) Quantification of DNA content profiles for time courses of cultures grown in different media concentrations (all 1:1000 initial dilution of a saturated culture). A) 1x Marine Broth (MB) B) 1/2x MB, C) 1/4x MB, and D) 1/16x MB. E) quantification of mean DNA content per time point (expressed as log2 of geometric mean). F) Nuclear doubling time, calculated by linear regression of data in E (timepoints from 0h to 36h). Error bars represent standard error of the slope. MB: Marine Broth media.

### Cell size

Next, we studied cell growth and the potential parameters that may be affecting it. First, we analyzed the effect of nutrient concentration on cell growth volume. We had initially observed that filtering saturated cultures through an 8-micron filter allows small uninucleated and binucleated cells to pass, while filtering cultures at 72 hours after inoculation (which also contain a large fraction of small cells, figure 2B) does not let any cells through (data not shown). This led us to hypothesize that nutrient concentration might affect the cell size of newborn cells.

To this end, we used FACS sorting to isolate fixed and stained cells from cultures at 60 hours after dilution into either 1x or 1/2x media (supplementary figure 3), and measured the cell volume by Coulter counter. We found that both for 2C (newborn cells) and 32C cells (coenocytes), the cell volume was significantly bigger in cells grown in 1x media than in 0.5x media (figure 4A).

**Figure 4:**
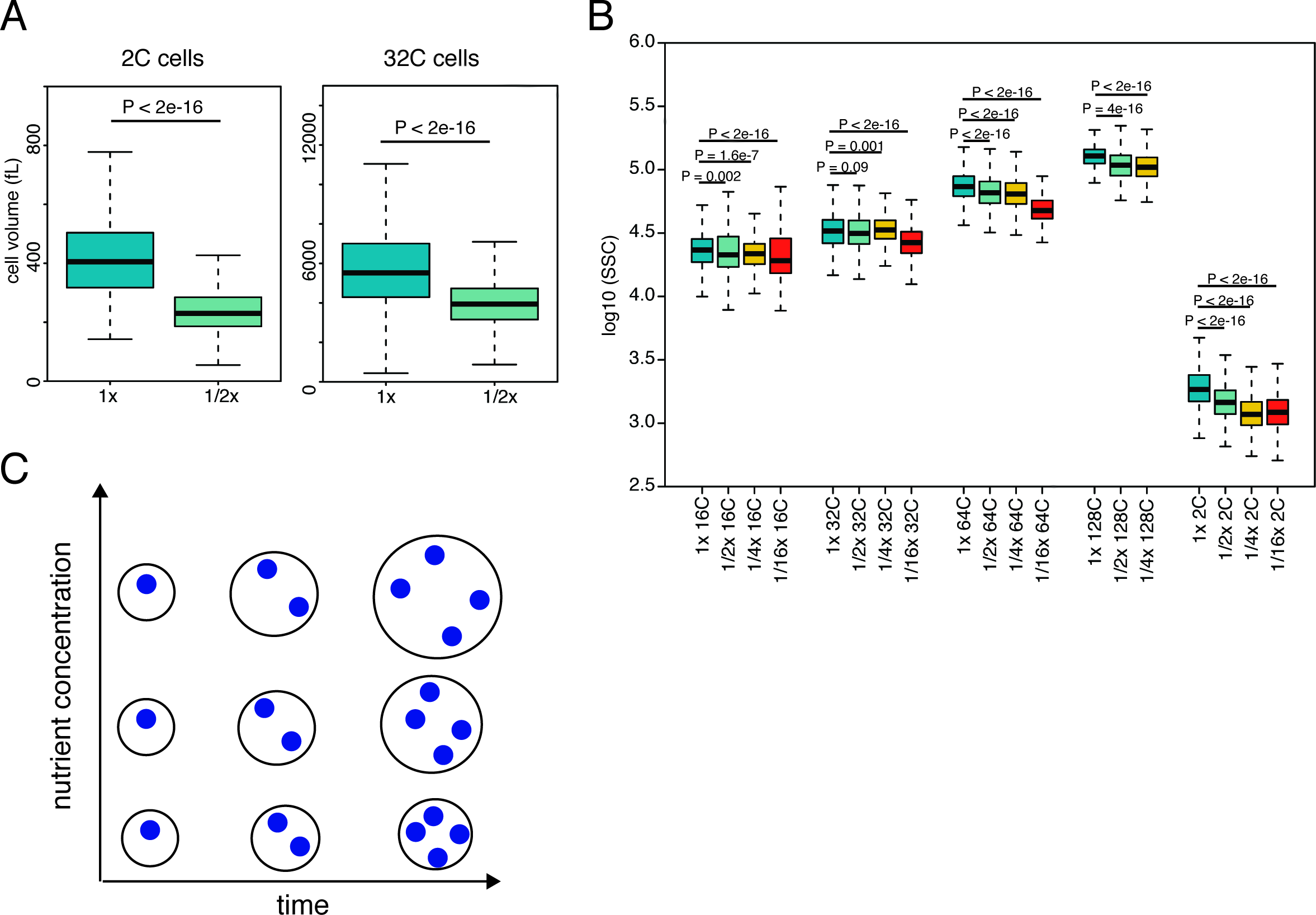
A) Comparison of cell volume measurement of 2C and 32C cells grown in either 1x or 1/2x nutrient concentration for 60 hours. Cells were isolated by FACS sorting based on DNA content and the volume was measured by Coulter counter. P values were computed using the Wilcoxon test. B) Comparison of SSC signal measured by flow cytometry between cells grown in 1x, 1/2x, 1/4x and 1/16x media concentration. Cells of each DNA content bin were compared from the time point when the population was most abundant (16C and 32C from 36h, 64C and 128C from 48h, and newborn 2C from 60h time points). P values were computed using the Wilcoxon test. In A and B, outliers were removed for visual clarity but were included in statistical analyses. C) a schematic representation of the relationship between nuclear division cycles and cytoplasmic volume growth of *S. arctica* in various conditions. In lower nutrient concentration, the timing of nuclear cycles remains the same, but the cell volume grows more slowly.

Next, we wanted to further examine the growth in cell volume throughout the cycle by a more dense sampling of the coenocytic cycle stages and under more media conditions. Forward scatter (FSC) and side scatter (SSC) measurements in flow cytometry generally provide information about cell size and cell shape (Tzur, Moore, Jorgensen, Shapiro, & Kirschner, 2011). Flow cytometry analysis of the sample containing a mixture of cells from different stages revealed that for *S. arctica* cells, SSC signal captures more variance than FSC signal (supplementary figure 1A). Having demonstrated the decrease of cell volume in lower nutrient concentration by direct measurement (see above), we used SSC signal as a proxy for cell volume across the time course experiment for multiple nutrient concentrations. We compared the SSC signal for cells of each nuclear content bin from experiment described in figure 3 at the time point where the population was most abundant (figure 4B). Although no significant difference was observed, presumably due to lower sensitivity of this measurement compared to Coulter counter, for cells grown in media concentrations of 1x to 1/4x for young coenocytes (up to 32C), coenocytes grown at 1/16x media concentration were significantly smaller in cell size (figure 4B). However, mature coenocytes (64C and 128C DNA content) showed a statistically significant trend in cell size with decreasing concentration of nutrients. In addition, the same trend was observed with newborn 2C cells (figure 4B). Taken together, these results indicate that the rate of growth of the cell volume decreases in lower media concentration; cells at lower nutrient concentration grow in volume more slowly, and give rise to smaller newborn cells.

Overall, our results show that the nuclear division cycles in the multinucleate cells of the ichthyosporean *Sphaeroforma arctica* occur synchronously and are governed by a timer mechanism that ensures constant and invariable timing of nuclear divisions, independent on nutrient concentration and independent of the growth rate of cell volume. The regulation of the cell size and the cell cycle of *S. arctica* is therefore distinct from the regulation of growth in filamentous fungi, where nuclear divisions within the coenocyte are autonomous (Anderson et al., 2013; Dundon et al., 2016), or the multiple fission cycle of the alga *C. reinhardtii*, and is reminiscent of the cell cycle in early animal embryos, in particular the insect preblastoderm. The uniqueness of these cell cycle features, the experimental tractability due to its simple life cycle in laboratory conditions, and evolutionary relevance due to its phylogenetic placement suggest *S. arctica* an interesting novel “non-model model” system for the studies of the cell division.

## Materials and methods

### Culture conditions and staining

*Sphaeroforma arctica* (strain JP6010) cultures were maintained in marine broth (Difco, 37.4 g/L) at 12°C. For media composition experiments, marine broth was diluted to desired concentration with artificial seawater (Instant Ocean, Aquarium Systems, 36 g/L). Cells were fixed in 4% paraformaldehyde in PBS for 15 minutes at room temperature, washed once with marine PBS (PBS with 35 g/L NaCl), and stained with DAPI (final concentration 5 ug/mL) in marine PBS. For microscopy, cells were washed twice with marine PBS before applying to the slide. For flow cytometry and cell sorting, samples were analyzed directly without any washing steps.

### Microscopy

Images were taken with the Zeiss Axio Observer Z.1 Epifluorescence inverted microscope equipped with Colibri LED illumination system and Axiocam 503 mono camera, using a 63x objective. Image processing was done with Fiji. For the DAPI channel, z-stacks of the images were combined using Z project function (maximum intensity).

### Flow cytometry and FACS sorting

Samples were analyzed using an LSRII flow cytometer (BD Biosciences, USA) and the data were collected with FACSDiva software. The data were then analyzed using FloJo and custom R scripts (available upon request). A mixed sample containing cells of all sizes and DNA contents was used to calibrate the measurements. SSC-A and FSC-A signals were used to detect population of cells. DNA content (DAPI) was detected using a 355nm laser with the 505nm longpass and 530/30nm bandpass filters. SSC-A signal was also used as a measure for cell size. Around 5,000 events were recorded in each measurement.

Cell sorting was performed using BD Influx cell sorter (BD Biosciences, USA). Sorting gates were set according to DAPI fluorescence, which was detected using a 355nm laser with the 400nm longpass and 460/50nm bandpass filters. Approximately 100,000 cells of each population were sorted into PBS with 35 g/L NaCl.

### Flow cytometry data analysis

Gating of subpopulations and subpopulation statistics was performed using FlowJo (Ashland, OR), and the data was further processed using custom R scripts (R core team, 2014)). log2 of geometric mean of DNA content was calculated as:

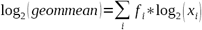

where f_i_ is the fraction of cells and xi the DNA content (ploidy) of each i-th DNA content bin. The rate of nuclear division was computed as linear regression of log_2_ of geometric mean of DNA content versus time. Statistical tests were implemented in R.

### Cell volume measurements using Coulter counter

Cell volume measurements were performed using the Coulter Counter Z2 (Beckman Coulter, USA). The data were collected using the Accucomp software. Blank sample (buffer) was used for background subtraction. Statistical tests were implemented in R.

## Acknowledgements

We thank Gautam Dey, Omaya Dudin, Michelle Leger, Sebastián Najle and Aaron New for discussions and comments on the manuscript, and Meritxell Antó for technical support. This work was supported by a European Research Council Consolidator Grant (ERC-2012-Co -616960) grant, and a grant (BFU2014-57779-P) from Ministerio de Economía y Competitividad (MINECO) to I.R.-T; the latest co-funded by the European Regional Development Fund (fondos FEDER). We also acknowledge financial support from Secretaria d’Universitats i Recerca del Departament d’Economia i Coneixement de la Generalitat de Catalunya (Project 2014 SGR 619).

---

**Supplementary figure 1:**
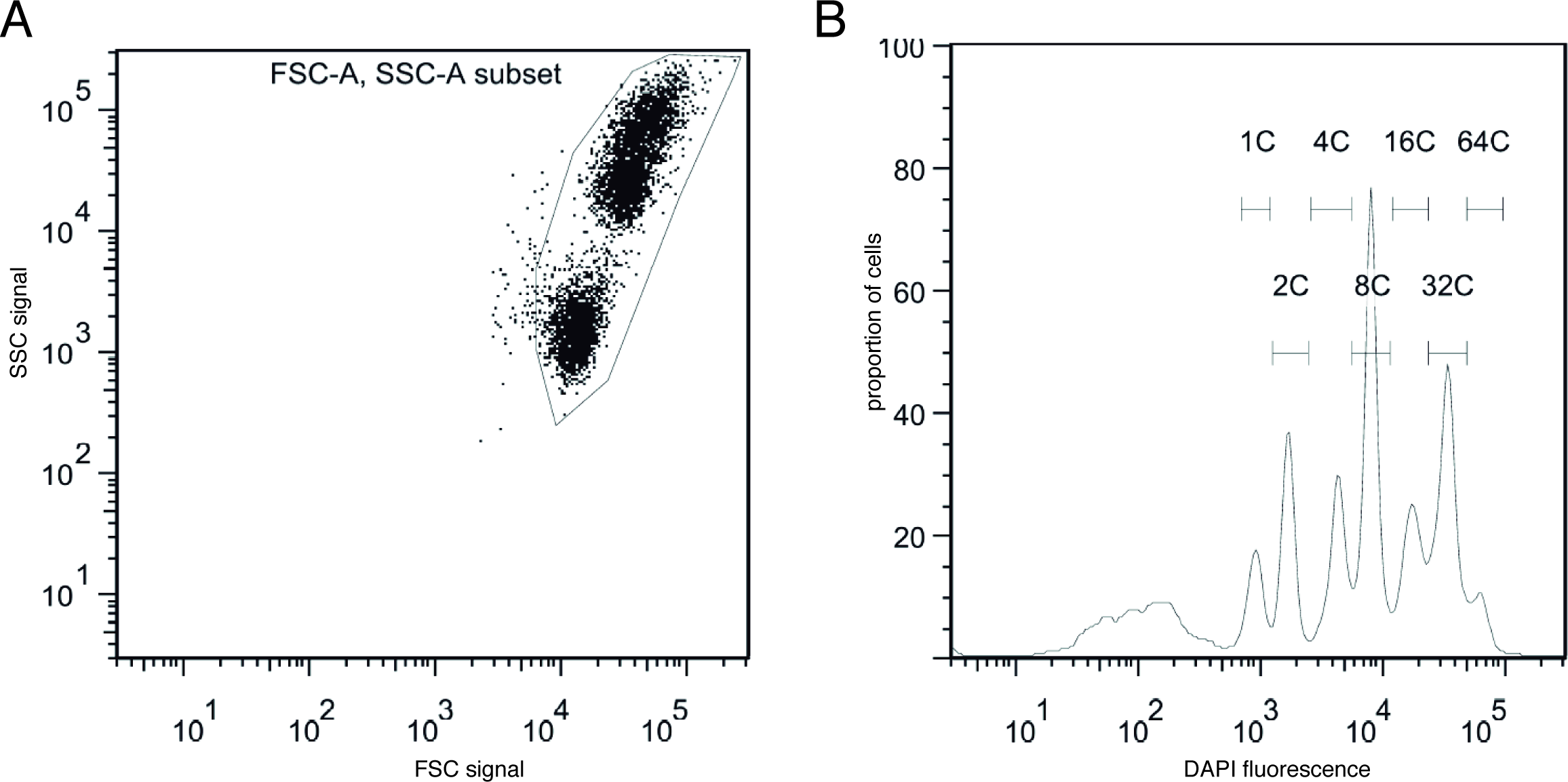
Flow cytometry profile of a mixed sample of 24h, 48h and 168h (mixed in 100: 100: 1 ratio) time points from the 1x MB time course. A) FSC-SSC profile. The gate shown was applied to the samples to discriminate cells from debris. B) DNA content. The DNA content bin gates were applied to the samples to quantify the fraction of cells at each DNA content.

**Supplementary figure 2:**
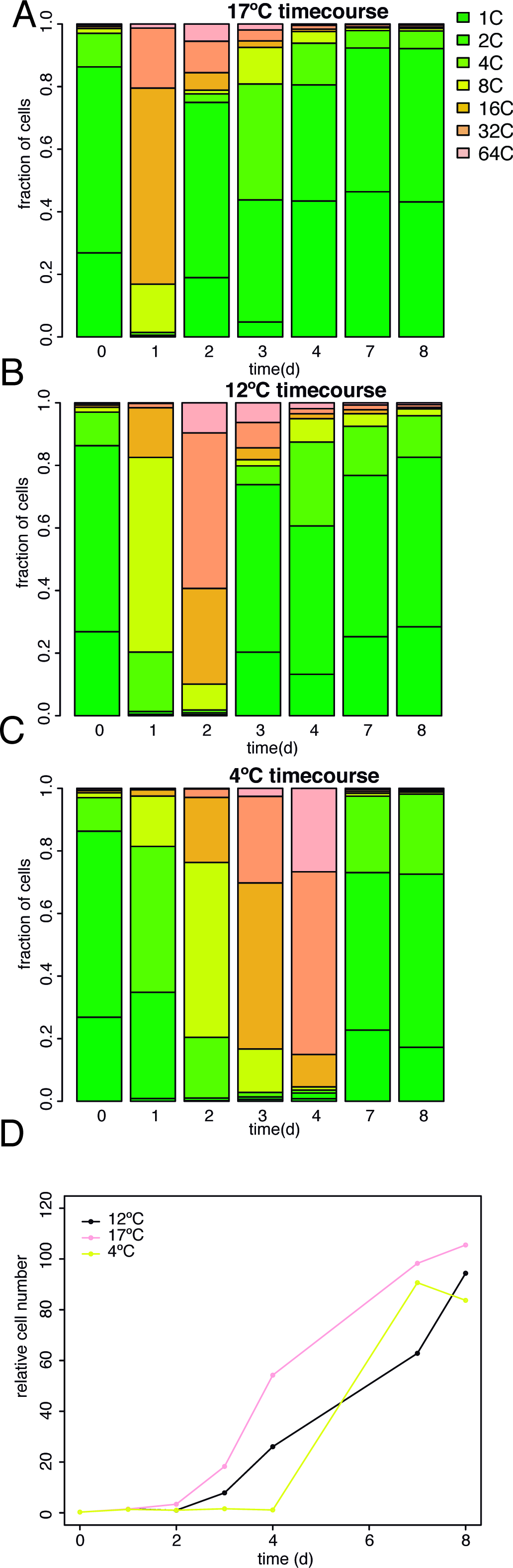
A-C) Quantification of DNA content profiles for cultures grown at different temperatures in 1x MB and 1:100 initial culture dilution. A) 17°C, B) 12°C, C) 4°C. D) Growth curves (relative cell density) for the conditions in A-C.

**Supplementary figure 3:**
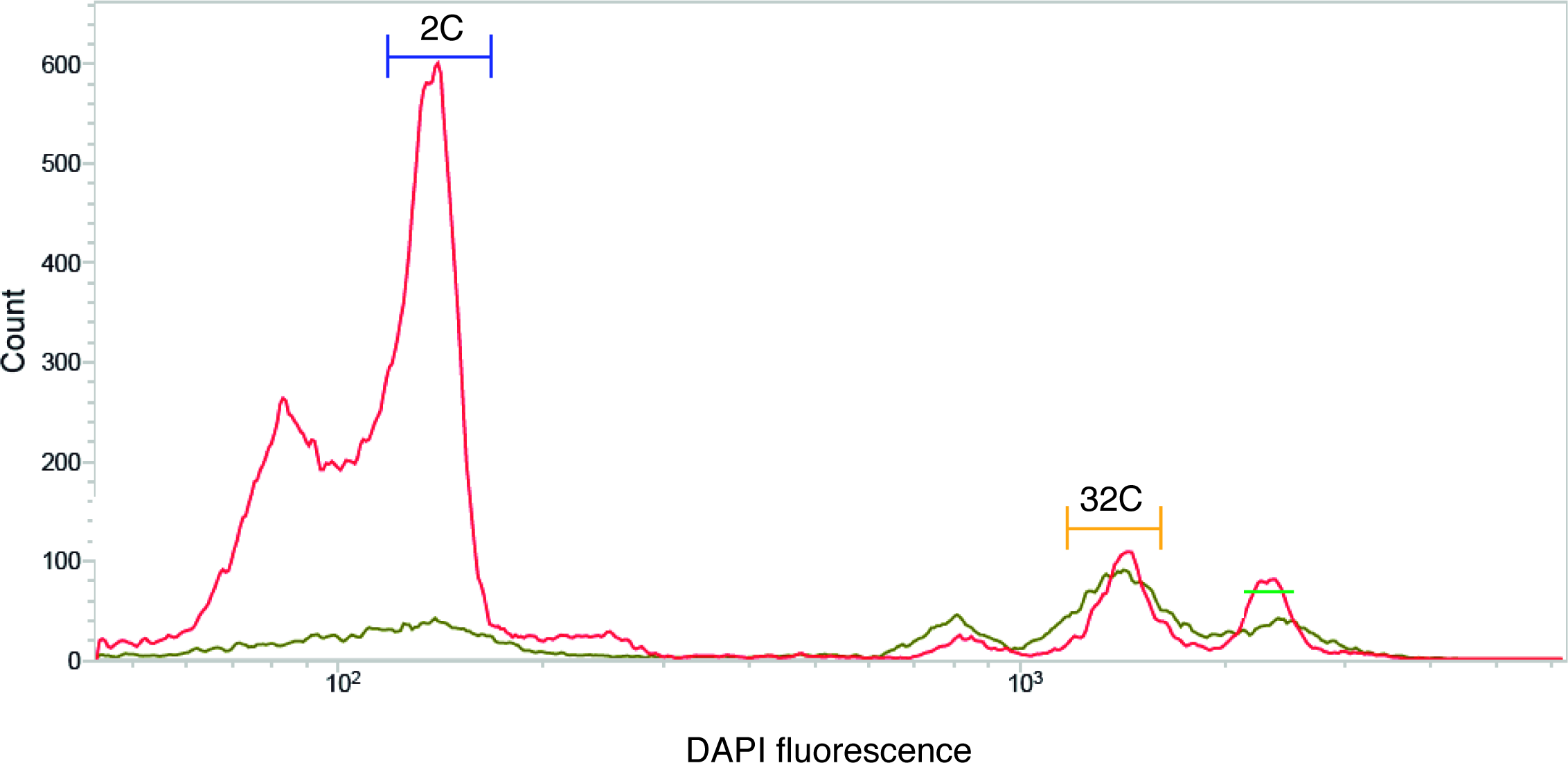
DNA content profiles used for cell sorting of cells with 32C and 2C DNA content from cultures grown in 1x MB (green line) and 0.5x MB (red line). The sorting gates are indicated on the plot.

